# Sustained Modulation of Deep Brain Circuits with Focused Ultrasonic Waves

**DOI:** 10.1101/2022.11.14.516484

**Authors:** Taylor D. Webb, Matthew G. Wilson, Henrik Odeen, Jan Kubanek

**Affiliations:** Department of Biomedical Engineering, University of Utah, 36 South Wasatch Dr, Salt Lake City, UT 84112; Department of Radiology, University of Utah, 50 2030 E, Salt Lake City, UT 84132

**Keywords:** noninvasive, deep brain, stimulation, ultrasound, durable

## Abstract

Transcranial focused ultrasound has the potential to noninvasively and systematically modulate deep brain circuits and impart sustained, neuroplastic effects in awake subjects. The intersection of these properties is critical for effective treatments of brain disorders, yet remains to be shown. Harnessing the full potential of transcranial ultrasound, we delivered 30-second stimuli into deep brain targets (left/right lateral geniculate nucleus) of non-human primates while they performed a visual discrimination task. This brief stimulation induced sustained and target-specific behavioral preference that persisted up to 15 minutes following the ultrasound offset. The polarity of the behavioral and neural effects suggested that ultrasound excited the stimulated circuits. The ultrasound was delivered into the deep brain daily for a period of more than 6 months, which enabled us to evaluate the safety of longterm stimulation. There were no detrimental effects on the animals’ discrimination accuracy over the course of this stimulation regimen. This study demonstrates ultrasound’s capacity to condition deep brain circuits in a safe and treatment-relevant manner in awake subjects, and provides a basis for effective and safe translations into humans.

**Highlights:**

- Transcranial ultrasound induces effective and sustained modulation of deep brain circuits.
- The deep brain modulation biases choice behavior of non-human primates.
- The deep brain modulation produces sustained elevation of high gamma activity.
- The stimulation, applied daily for several months, is safe.

## Introduction

Mental and neurological disorders are frequently resistant to existing treatments [1–8]. Targeted neuromodulation has provided treatment options for some of the patients but existing approaches—such as deep brain stimulation, electroconvulsive therapy, or transcranial magnetic stimulation—are either invasive or lack the capacity to directly modulate specific deep brain regions. These limitations have yielded variable response, with only a subset of patients responding to the stimulation.

Transcranial low-intensity focused ultrasound can noninvasively modulate deep brain circuits with millimeter precision [9]. The effects on the target circuits can be either transient [10–18] or durable [19–26], depending on the stimulus duration. While transient perturbations are useful for diagnostic and guidance applications, successful therapeutic translations require the capacity to induce sustained changes in behavior following the modulation of specific deep brain circuits (Fig. 1A).

**Figure 1:**
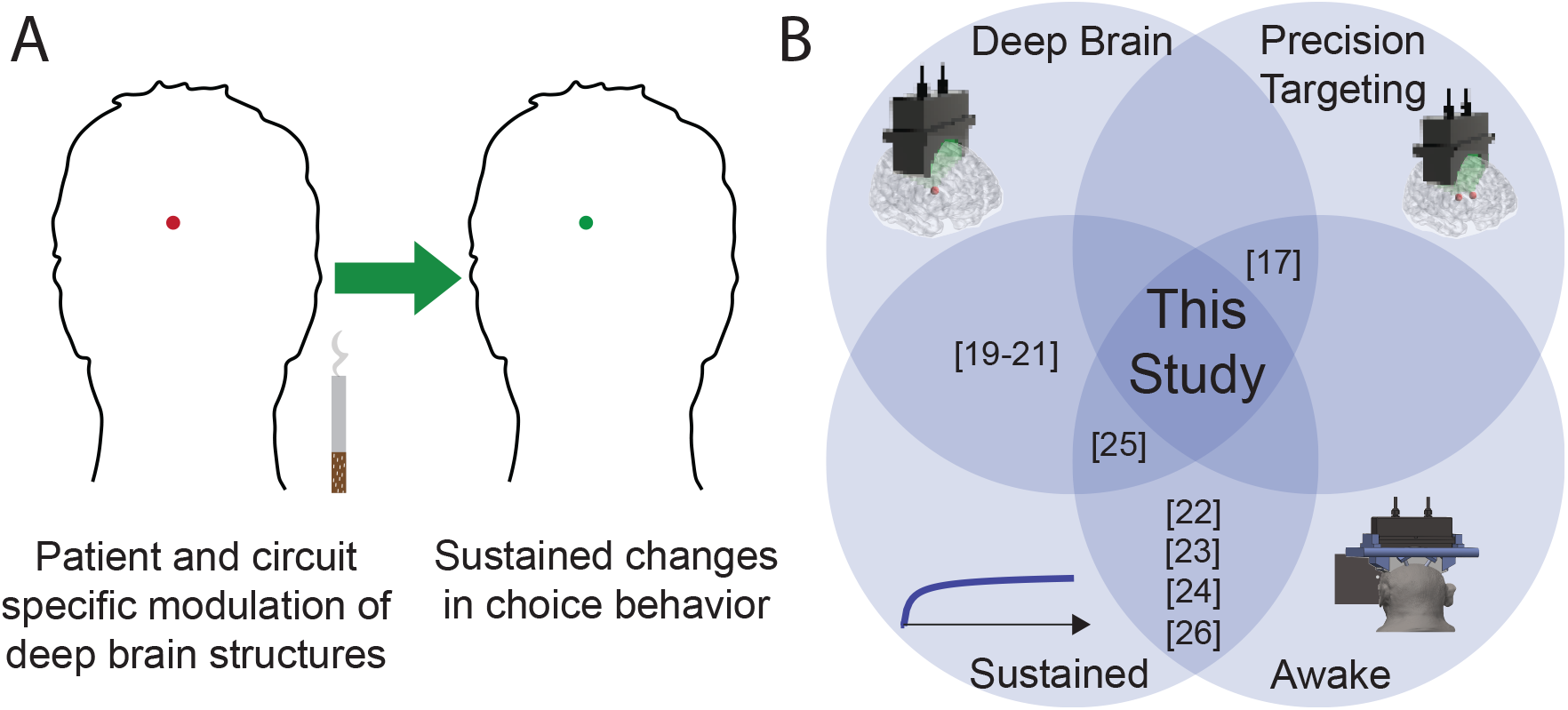
Non-Invasive Precision Medicine for the Brain. **(A) Durable Precision Therapies:** Ultrasound uniquely combines the capacity to noninvasively target deep brain circuits with high precision and flexibility. Its potential for inducing sustained changes in the targeted circuit is a key step towards large-scale treatments of nervous system disorders (e.g., nicotine dependence). **(B) The Key Strengths of Ultrasound:** Transcranial focused ultrasound has the capacity to modulate deep brain targets, produce sustained effects on the circuits, target brain circuits with high precision, and mediate effects while applied in awake subjects. This study combines these strengths. The brackets provide references to studies within the respective domains.

Thus far, studies showing durable changes in target circuits [19–26] have either required anesthesia or only used single-element transducers, which have limited spatial resolution and targeting flexibility. The present study now addresses these shortcomings, demonstrating precise targeting of multiple deep brain regions to induce sustained and region-specific neuroplastic effects in non-anesthetized subjects (Fig. 1B). Together with confirming its safety, the study brings the approach closer to therapeutic options for patients suffering from treatment-resistant disorders of brain function.

## Results

### Noninvasive, Sustained, and Reversible Neuromodulation

This study leverages Remus [17], a phased array system that programmatically delivers ultrasound into specific deep brain targets (Fig. 2A). We engaged the subjects in an established visual discrimination task [14, 17, 27, 28] that enabled us to evaluate the polarity, magnitude, and duration of the neuromodulatory effects. In this task, two targets appear on the left and right part of the screen, with controllable delay between their onsets. The subject’s task is to look at the target that appeared first (Fig. 2B). Using this system, we targeted the left and right lateral geniculate nucleus (LGN; Fig. 2C). Inhibition/excitation of the LGN during this task is known to result in an ipsilateral/contralateral bias in subjects’ choice behavior [14, 27–30]. The phased array system enables flexible and reproducible targeting of the left and right LGN without any changes to the experimental setup [17], which allowed us to systematically evaluate lateralized effects and the long-term safety of the stimulation. We found that a single 30-second stimulation of the LGN induced a sustained but reversible contralateral bias in the subjects’ visual choice behavior (Fig. 2D). We quantified the effect in the same way as in a previous study [14] and tracked the ultrasound-induced changes in the choice preference as a function of time throughout the session (Fig. 2E). The figure shows that the effect was specific to the stimulated hemisphere. In particular, stimulation of the left/right LGN induced a rightward/leftward bias in the subjects’ choices. This effect polarity obtained in this task suggests that ultrasound excited the stimulated circuits [14]. The effects outlived the stimulation by up to 15 minutes (final time point for which the effect in Figure 2d is significant; two-tailed t-test, *p <* 0.05).

**Figure 2:**
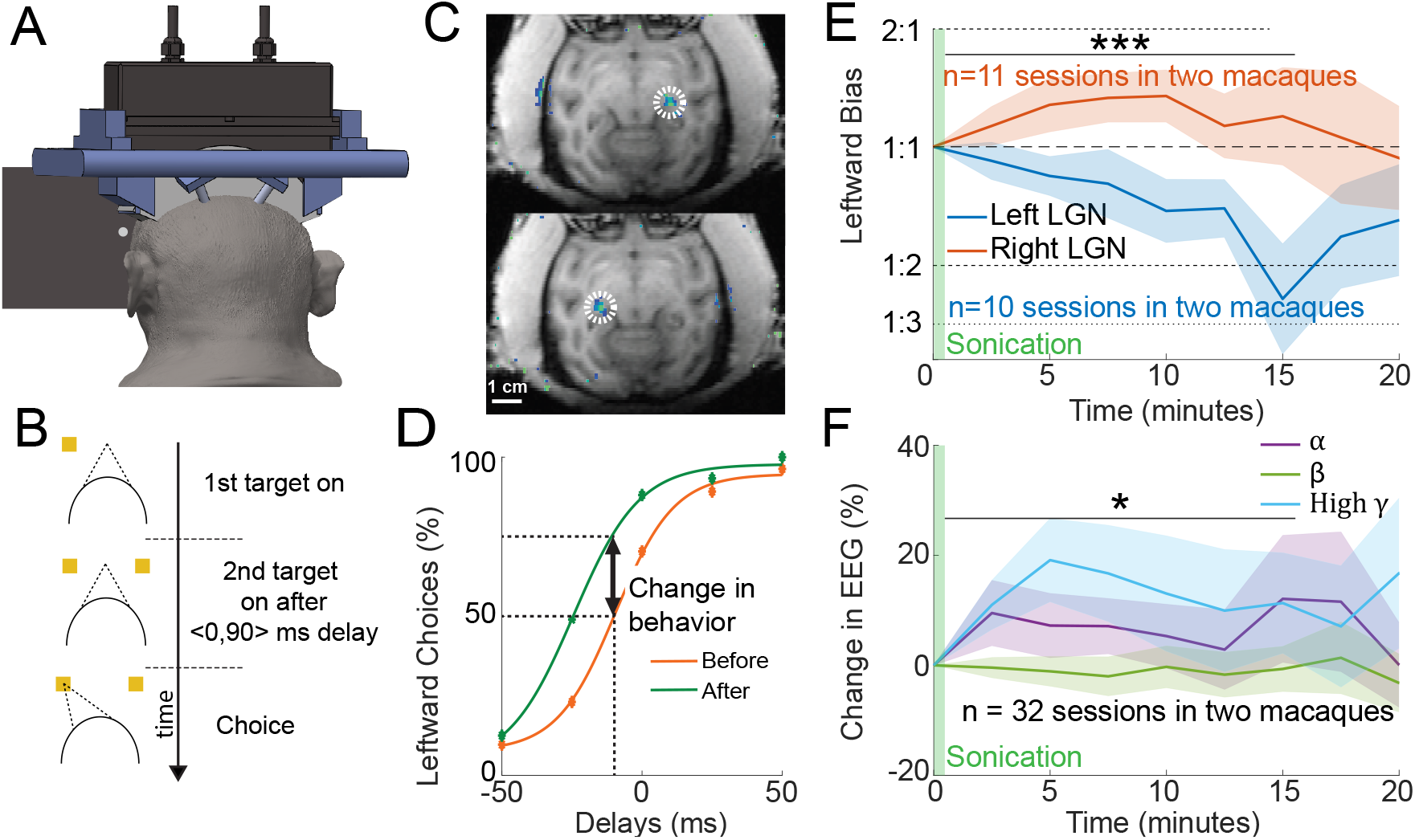
Noninvasive, Sustained, and Reversible Modulation of Deep Brain Circuits and Choice Behavior. **(A) Apparatus:** A system for reproducible delivery of ultrasound into deep brain targets [17] delivered 30-s of ultrasonic stimulation into to the left/right LGN while subjects performed a visual discrimination task. **(B) Choice task:** Following fixation of a central target, a target appears in the right or left hemifield, followed by a second target in the opposite hemifield. There is a random, 0-90 ms delay between the onset of the two targets. The subject is rewarded for selecting the target that appeared first. **(C) Deep Brain Targeting:** Validation of the left/right LGN targeting [17]. This system enables fully programmatic delivery of ultrasound into specified deep brain targets; no translation of the transducer is necessary. **(D) Quantification of the Effects on Choice Behavior:** Example session of right LGN stimulation, demonstrating a strong contralateral effect on choice behavior. A baseline epoch prior to the ultrasound (orange) is used to establish the point of equal preference for each session. In all subsequent plots, we quantify the proportion of leftward choices at this point of equal preference following the stimulation (green). **(E) Sustained Modulation of Behavior:** Sonication of the right (red) or left (blue) LGN induces a persistent contralateral bias in the animals’ choices. The data are shown as mean *±* s.e.m. These data only include early sessions before animals could adapt to the stimulation (see text for details). The result over all datasets is shown in Fig. 3. : \*\*\**p <* 0.001: A two-way ANOVA detected a significant effect of the stimulated side on the choice behavior (see text for details). **(F) Sustained Increase in Neural Activity:** Neural activity was recorded from parieto-occipital leads (see panel a). Mean *±*s.e.m. alpha, beta, and high gamma activity as a function of time. The modulation by the frequency band was assessed using two-way ANOVA (see text for detail). *: *p <* 0.05.

We quantified the effects on choice behavior using a two-way ANOVA, with factors sonicated side and time. Notably, in this analysis, we only included times following the ultrasound offset. This way, the reported effects are free of artifacts that can be associated with transcranial ultrasound [31, 32]. The effect of the sonicated side was highly significant (*F* (1, 45) = 17.87, *p* = 1.1 *×* 10^−4^). Time or its interaction with side were non-significant.

These effects on choice behavior were accompanied by an increase in high gamma activity recorded over visual cortex (Fig. 2F). As with the behavioral analysis, we only assessed statistical significance of effects after the ultrasound offset, to make sure the recordings were not affected by a potential artifact that could be associated with the ultrasound. A two-way ANOVA, with factors signal frequency and time, identified a significant effect of signal frequency (*F* (2, 156) = 3.3, *p* = 0.040). Time or its interaction with frequency were non-significant. Since high gamma activity is tightly correlated with multi-unit spiking [33–35], this effect suggests an excitation of the stimulated circuits.

The behavioral and neural effects showed comparable temporal profile (Fig. 2D-E). Moreover, both effects suggest a neural excitation of the stimulated circuits [14].

### Adaptation

The stimulation was applied to the subjects over the course of many months. As a consequence, the monkeys learned to compensate for the stimulatory effects. A compensation is expected because an ultrasound-induced bias decreases the discrimination accuracy and therefore the reward income for the animals.

The inclusion of all sessions in the analyses (Fig. 3A) preserves the effect (*F* (1, 186) = 8.8, *p* = 0.0033) but reduced its magnitude compared to the results obtained within the first and sixth month of stimulation (Fig. 2E). Fig. 3B shows the average effect across the progressive months of stimulation, measured in the 10-15 minute time window following each day’s stimulation. The figure shows that the effect was strong during the first month of the stimulation—the animals’ decisions were biased by the ultrasound by more than 2:1, compared to the default balanced proportion before ultrasound was applied. This prominent effect gradually declined thereafter—the animals became less biased by the stimulation and became more accurate in the task. Given this result, at month 5, the relatively high-energy sonication regimen was paused for several weeks in each animal. Resuming the stimulation, the effect and its adaptation were replicated (Fig. 3B).

**Figure 3:**
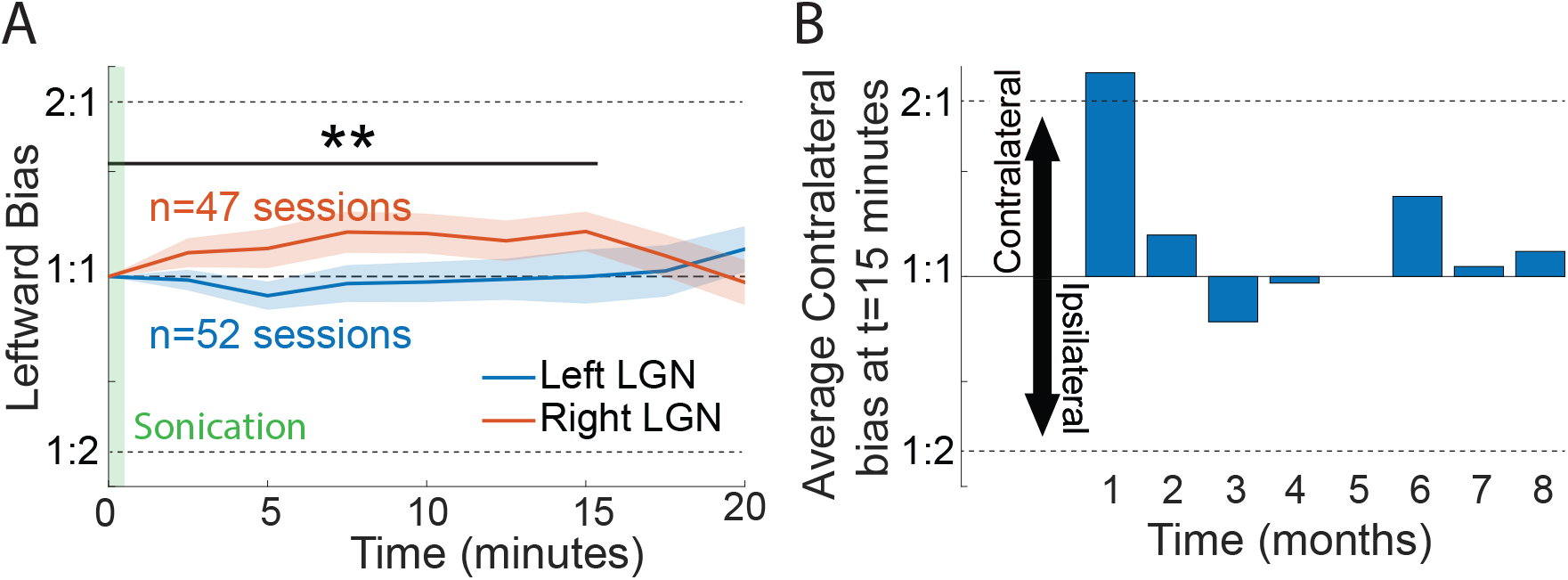
Behavioral Adaptation. **(A)** Same format as in Fig. 2E, for data of all recorded sessions. **(B)** The effect is prominent within the first month of stimulation and gradually declines thereafter. See text for details.

### Safety

The reproducible daily stimulation of the LGN over the course of many months provides crucial information on the safety of sustained ultrasonic neuromodulation in the primate brain. The task used in this study has been used in neurology for more than a century as a sensitive readout of the impact of stroke or other perturbations of visual regions [17, 27, 28, 36]. Specifically, damage to the LGN would manifest as a decrease in the subject’s visual discrimination accuracy [17, 27, 28, 36]. No such decrease was observed over the months-long stimulation regimen (Fig. 4). The subjects showed either stable or improved accuracy over time (the improvement in the second set of sessions for subject B is significant (*r* = 0.80, *p <* 0.001); all other linear fits were non-significant). Thus, the subject’s behavior in this sensitive task reveals no evidence of damage as a result of chronic ultrasonic stimulation.

**Figure 4:**
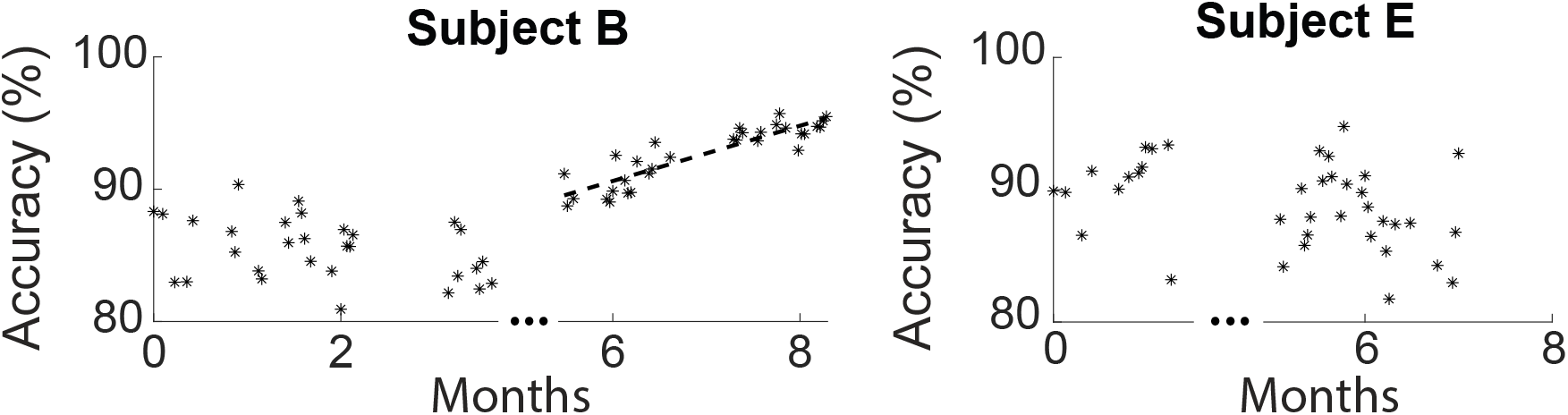
Chronic Modulation of Primate Deep Brain Circuits with Ultrasound Is Safe. Repeated application of neuromodulatory ultrasound to deep brain circuits of non-human primates is safe. Significant damage to the LGN in this sensitive visual discrimination task would cause a notable decline in subjects’ visual discrimination performance. We observe no such decline, with both subjects showing consistent or improving performance across sessions. The time of the pause in the stimulation regimen is indicated by the ellipsis.

### Stimulation Parameters

As a part of this chronic stimulation study, we aimed to systematically investigate the effects of individual stimulation parameters (Tab. 1 and Tab. 2). Unfortunately, the observed adaptation to the stimulation (Fig. 3) reduced the power of this analysis, and so any claims on relative advantages of some parameters over others would be inconclusive.

**Table 1:** Sonication Parameters in Subject B. Left to right columns: Spatial peak temporal average intensity (*I*_SPTA_), spatial peak temporal peak intensity (*I*_SPPA_), duty cycle (DC), the number of sessions recorded during sonication of each LGN, the number of sham sonications, and whether or not the beam was scanned throughout the LGN (see Methods details). Intensities are given in *W/cm*^2^.

| $I_{\text{SPTA}}$ | $I_{\text{SPPA}}$ | DC (%) | Left | Right | Sham | Scan |
| --- | --- | --- | --- | --- | --- | --- |
| 1 | 30 | 3.6 | 8 | 6 | 0 | No |
| 1 | 30 | 3.6 | 8 | 7 | 0 | Yes |
| 1 | 10 | 14.4 | 8 | 8 | 8 | Yes |
| 2 | 60 | 3.6 | 8 | 8 | 0 | Yes |

**Table 2:** Sonication Parameters in Subject E. Same format as in Table 1.

| $I_{\text{SPTA}}$ | $I_{\text{SPPA}}$ | DC (%) | Left | Right | Sham | Scan |
| --- | --- | --- | --- | --- | --- | --- |
| 1 | 60 | 1.8 | 9 | 8 | 0 | Yes |
| 2 | 15 | 14.4 | 4 | 5 | 0 | Yes |
| 2 | 60 | 3.6 | 7 | 5 | 5 | No |

### Active Sham Control

Thus far, there are two controls for potential generic artifacts associated with ultrasound [31, 32]. First, the effects outlive the stimulation by up to 15 minutes (Fig. 2E). Second, the delivery of focused ultrasound into the left and the right LGN elicited effects of opposite polarities (Fig. 2E). We included a third control in the form of active sham. The sham sonication delivered into the brain the same amount of acoustic energy but the phases applied to each element of the transducer were set at random, thus creating an unfocused beam. Fig. 5 shows the behavioral effects during this active sham. There were no significant changes in the choice behavior in this control condition.

**Figure 5:**
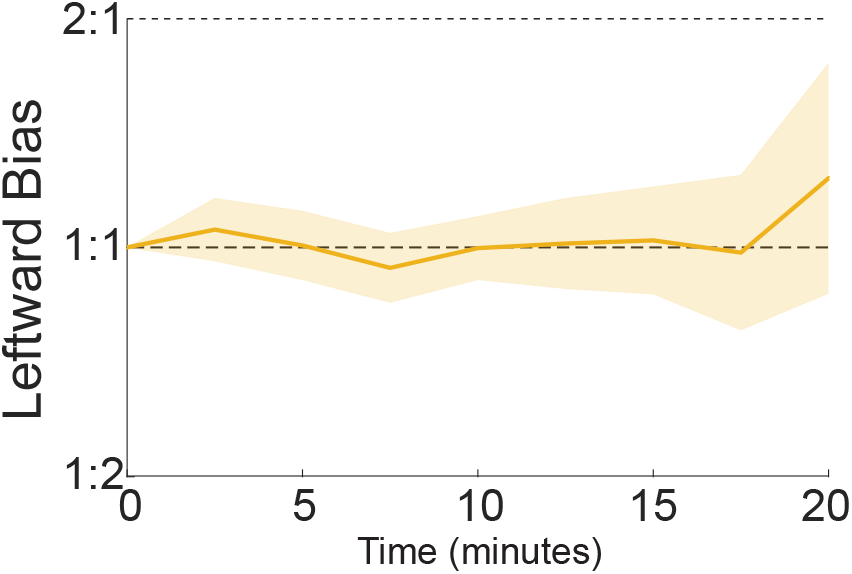
Active Sham. Same format as in Fig. 3A. Sham sonications delivered into the brain the same amount of energy but in an unfocused manner.

## Discussion

This study demonstrates that transcranial focused ultrasound can effectively and safely stimulate deep brain circuits in primates. Just 30 seconds of low-intensity ultrasound produced 15-minute long effects on the subjects’ choice behavior and high gamma activity in visual cortex. Therefore, low-intensity ultrasound is capable of inducing durable stimulatory effects within confined deep brain circuits of primates. Effective and safe treatments of brain disorders require an approach that modulates deep brain targets, does so precisely, in a sustained manner, and in subjects who are not anesthetized. This study is unique in that it combines all four strengths (Fig. 1B). We have modulated two deep brain targets, the left and the right LGN. These targets were modulated at high spatial precision using a phased array (Fig. 1C). We have demonstrated that the effects of brief stimulation trains outlast the stimulation—a key premise for inducing a durable reset of malfunctioning circuits. And finally, the stimulation was performed in primates who were not anesthetized. Anesthesia should be avoided as it profoundly impacts cortical excitability, including the excitability by ultrasound [37]. Moreover, anesthesia should be avoided to maximize the spectrum of patients who can receive treatment.

The finding that relatively short trains of stimulation (30 seconds used here) induce durable effects within the target circuits align with previous findings, which report effects on NHP behavior [25], resting-state fMRI [19–21], and motor evoked potentials in humans [26]. The present study applied these protocol systematically to two distinct targets with predicable changes in behavior. This allowed us to assess the polarity of the sustained effects. Both the behavioral and neural results suggested that the ultrasound excited the stimulated circuits.

The result reported here is in line with a recent mechanistic study that recorded discharge activity of primary rat cortical neurons in response to low-intensity ultrasound [38]. A 40-s stimulation increased the neuronal excitability for up to 6 hours. The protocol used a neuronal culture and a lower ultrasound frequency (200 kHz as opposed to 480 kHz), but both studies demonstrate that ultrasound can excite neural tissues in a sustained manner.

We found that ultrasonic stimulation of the LGN evoked gamma activity over visual cortex, and, to a lesser extent, also increased alpha activity. Both effects have been observed in awake macaque monkeys following visual stimulation of the LGN [39]. These effects reflect specific directions of information flow between the LGN and the visual cortex during stimulation. The finding that visual and ultrasonic stimulation of the LGN evokes similar effects in the visual cortex provides additional evidence for effective engagement of the target. However, how exactly ultrasound mediated its excitatory effects should be investigated using invasive procedures in the future.

The duration of effects reported in this study is on par with the effects elicited by trains of TMS pulses of a comparable duration. [40] This is a notable finding given that TMS applied to the brain for about 40 minutes per day for several weeks can induce persistent improvements in major depression [41]. Ultrasound could be applied in a similar, repeated mode, with the key advantage that it can directly modulate diseased deep brain targets. This targeted action is expected to increase the effectiveness and safety of the treatments.

The sustained effects addresses the lingering concern that ultrasonic neuromodulation is associated with an auditory or vestibular artifact [31, 32]. We quantified the behavioral data in this study after the ultrasound was turned off. During that time, there was no stimulation and so there could be no auditory or vestibular confound. Moreover, the effects were stimulation-hemisphere-specific and no effects were observed during active sham stimulation.

This study informs on the safety of long-term ultrasound stimulation. The ultrasound was applied to the deep brain targets almost every working day over several months. Moreover, we delivered into the targets intensities higher than the FDA 510(k) Track 3 levels for diagnostic ultrasound imaging: *I*SPTA values of up to 2 W/cm^2^ in situ. Using a behavioral task that is often used in neurology to detect harm to nervous tissue [27, 28, 36], we found, if anything, improvements in the subject’s discrimination accuracy. Therefore, no safety concerns were detected during chronic stimulation of deep brain circuits in primates.

The study has two limitations. First, we have applied stimuli of limited duration (30 seconds). Future studies should test effects of stimuli of much longer duration, e.g., up to about the session length of standard applications of TMS treatments to depression (about 40 minutes). Second, our subjects adapted to the stimulation after about 1 month (Fig. 3B). This is expected given that the adaptation leads to increased reward income. As a consequence, we were unable to systematically test the effects of the individual stimulation parameters in this paradigm. Future studies should either apply the stimulation less frequently, or apply the stimulation to higher-level visual regions to avoid adaptation.

In summary, this study demonstrates effective and safe chronic modulation of deep brain circuits with transcranial ultrasound. The study highlights the full potential of the approach, modulating primate deep brain circuits in a sustained manner, systematically, and in non-anesthetized subjects. These results encourage ensuing applications of sustained modulation of deep brain circuits in humans.

## Methods

### Animals

Two male macaque monkeys (*macaca mulatta*, age 7 and 5 years, weight 9.5 and 12 kg) participated in this study. The procedures were approved by the Institutional Animal Care and Use Committee of the University of Utah.

### Task

We trained two NHPs to perform a visual discrimination task that is often used in neurology and neuroscience to quantify the magnitude and polarity of neuromodulatory interventions [14, 27, 28].

Each trial begins with the subject acquiring a central fixation target. After a random delay, a target appears in either the left or right visual hemifield. A second target then appears in the opposite hemifield after a delay between 0 and 90 ms. The subject was rewarded for looking at the target that appeared first. Subjects received a liquid reward with a probability 0.5 *<*= *p <*= 1 if they looked at the target that appeared first within 2 s. In the condition in which both targets appear at the same time both subjects are rewarded for either choice.

During the task, the subject was seated head-fixed in a comfortable primate chair while presented with the visual stimuli on a monitor. The subject’s eye movements were tracked and recorded using an infrared eye tracker (Eyelink, SR Research, Ottawa, Canada).

### Protocol

Sonication of the right and left LGN was interleaved across sessions. The left LGN was sonicated in 52 sessions (32 in subject B and 20 in subject E) and the right LGN in 47 sessions (29 in subject B and 18 in subject E). Changes in behavior were quantified by fitting a sigmoid to a subject’s average choice behavior in five minute blocks (Fig. 2C, negative delays represent the cases in which the target in the right hemifield appeared first). In each session, baseline behavior was measured in the five minutes before the sonication. In this time window, we establish the delay at which the subject had an equal preference for either target. Changes in the subject’s behavior in subsequent five-minute blocks were then quantified by measuring the change in the subject’s preference for the leftward target at the baseline delay.

The subjects completed a minimum of 600 trials to provide an adequate baseline before the 30-second sonication. The subjects generally worked daily throughout the work week, over a period of several months.

Both subjects were allowed to work ad libitum following the sonication. Subject B performed the task for an hour or more while Subject E’s performance was typically shorter. Thus, the amount of available data decreases with session time.

A total of 61 sessions in subject B and 38 sessions in subject E were collected. Sessions in which the subject worked for less than five minutes following the sonication were excluded. Occasionally, subjects would fall asleep, resulting in periods in which they did very few trials. Thus, five minute windows in which the subject worked at a rate of less than 1 trial every 10 seconds were also excluded. The 20 minute time point displayed in Fig. 2D contained 51 sessions for subject B and 20 sessions for Subject E.

### EEG recordings

EEG signals were recorded by attaching electrodes to the surgically implanted pins used to secure the transducer to the subject’s skull. A front pin was used as a reference electrode and the signal from the two rear pins was averaged to record the EEG signals. The signals were low-pass filtered at 7 kHz, sampled at 20 kHz, and notch-filtered at 60, 120, and 180 Hz.

Power in the *α* (7 to 12 Hz), *β* (12 to 30 Hz), and high *γ* (70 to 320 Hz) bands was measured using the fast Fourier transform within 500 ms windows. Windows in which the EEG signal exceeded 100 *μ*V were excluded. For each frequency band, the power over the 500 ms windows were averaged in 5 minute increments every 2.5 minutes (Fig. 2E). The baseline value was determined by averaging the five minute window prior to the sonication.

### Acoustic Parameters

All ultrasound stimuli had a center frequency of 480 kHz. Averaged across the two subjects, the estimated pressure at focus was 46% of the free field value (estimated using MR thermometry [17]). Thus, the values given in Tab. 1 and Tab. 2 assume that the *in-situ* intensity is de-rated by a factor of 21% relative to the free-field intensity. Each sonication lasted thirty seconds and consisted of a train of 30 ms pulses. The individual parameters used are provided in Tab. 1 and Tab. 2. The pulsing was achieved with a rectangular envelope.

### Beam Scanning

The focused, 480 kHz ultrasonic beam impacted only a portion of the LGN (Fig. 2C). To investigate the effect of partial modulation we electronically focused the beam into five adjacent locations. Tab. 1 and Tab. 2 give the parameters for which beam scanning was applied. Specifically, the original central target was complemented by targets *±*1.5 mm in the left/right dimension and *±*3 mm in the inferior/superior dimension. We found that this approach provided the same principal result as the original, single-target result.

### Statistical analyses

The two-way ANOVA, used to establish the difference in behavior between the two stimulation targets, was evaluated in time windows of [0.5,5.5], [5.5,10.5], and [10.5,15.5] minutes following ultrasound the offset of the stimulation. The windows start at 0.5 in order to exclude trials performed during the sonication.

## Acknowledgments

This work was funded by the NIH grants R00NS100986, RF1NS128569, and F32MH123019, by the Focused Ultrasound Foundation, and by the Margolis Foundation. We thank Caroline Garrett for veterinary assistance.

